## Supplementary Figures 1 and 2 for "The adenosine 2a receptor and TIM3 directly inhibit killing of tumor cells by cytotoxic T lymphocytes through interference with cytoskeletal polarization"

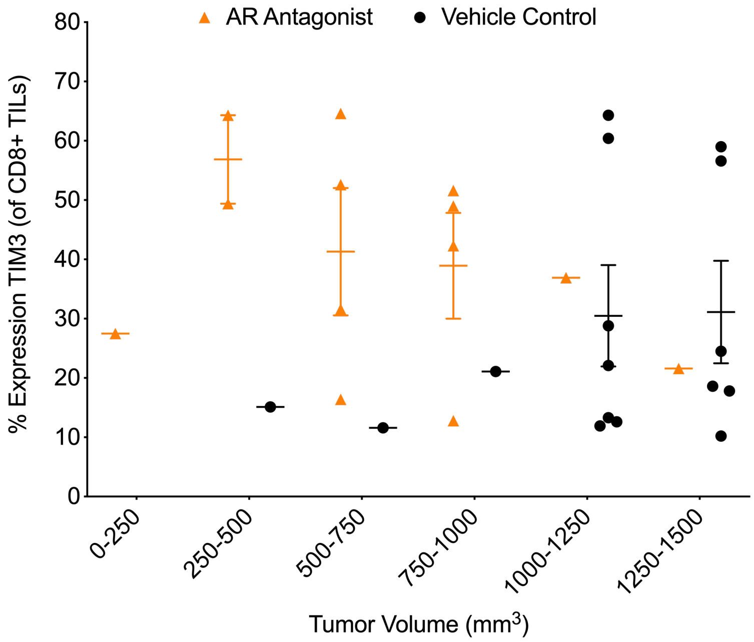


**Figure S1 – preferential upregulation of TIM3 upon A2aR agonist treatment on T cells from smaller tumors**

Percentage of CD8$^+$ TIL from RencaHA tumor-bearing BALB/c mice treated with or without the A2aR antagonist ZM 241385 at 10 mg/kg i.p. expressing TIM3 are given as a function of tumor size as mean ± SEM for N=6 experiments.

**A**

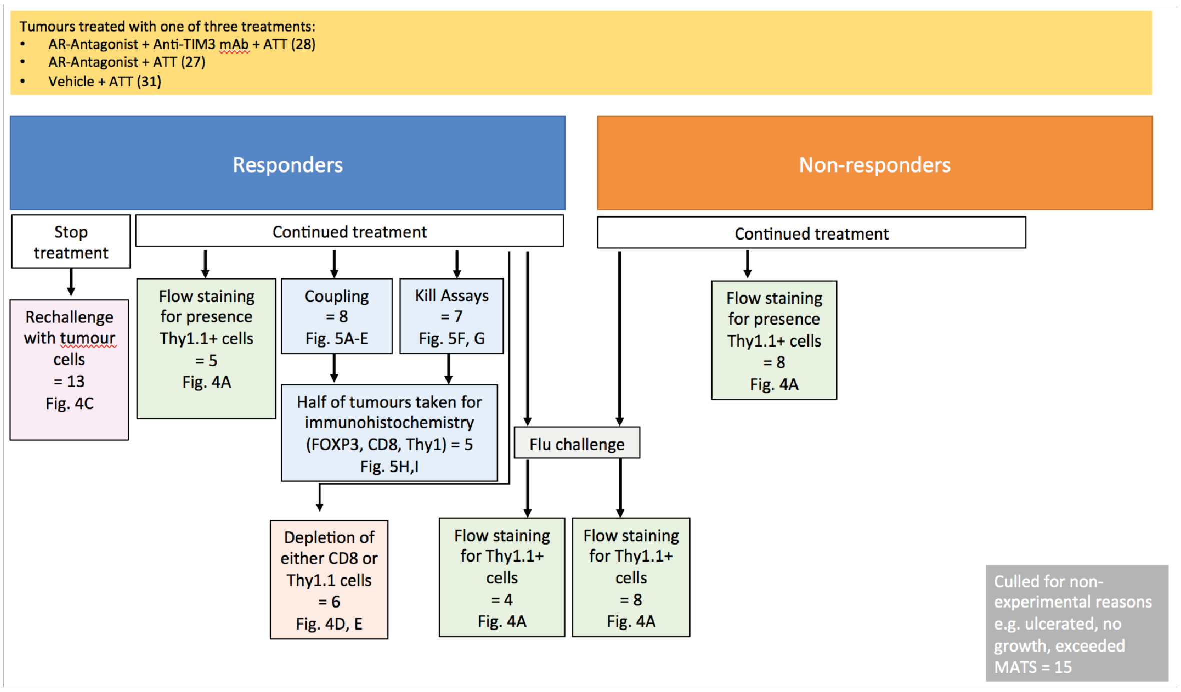


**B**

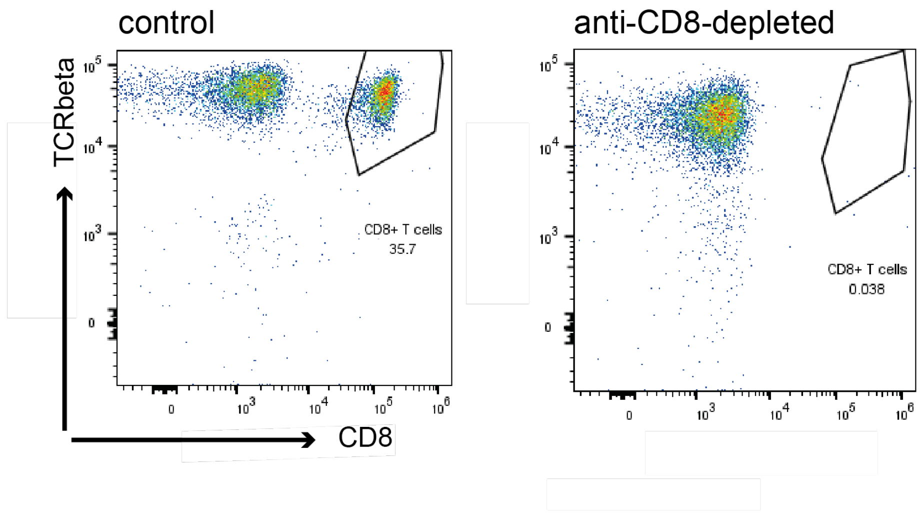


### Figure S2 – Supporting technical data for *in vivo* tumor growth experiments upon adoptive CTL transfer

**A** Numbers of mice used across the different parts of the experiments in Figs. 4 and 5 are given. **B** The efficiency of the depletion of CD8$^{+}$ T cells in Fig. 4E is shown in representative FACS plots from blood samples two days after the injection of the depleting antibody or control.
